# Nanobubbles are Non-Echogenic for Fundamental-Mode Contrast-Enhanced Ultrasound Imaging

**DOI:** 10.1101/2022.03.25.485890

**Authors:** John Z. Myers, J. Angel Navarro-Becerra, Mark A. Borden

## Abstract

Microbubbles (1–10 µm diameter) have been used as conventional ultrasound contrast agents (UCAs) for applications in contrast-enhanced ultrasound (CEUS) imaging. Nanobubbles (<1 µm diameter) have recently been proposed as potential extravascular UCAs that can extravasate from the leaky vasculature of tumors or sites of inflammation. However, the echogenicity of nanobubbles for CEUS remains controversial owing to prior studies that have shown very low ultrasound backscatter. We hypothesize that microbubble contamination in nanobubble formulations may explain the discrepancy. To test our hypothesis, we examined the size distributions of lipid-coated nanobubble and microbubble suspensions using multiple sizing techniques, examined their echogenicity in an agar phantom with fundamental-mode CEUS at 7 MHz and 330 kPa peak negative pressure, and interpreted our results with simulations of the modified Rayleigh-Plesset model. We found that nanobubble formulations contained a small contamination of microbubbles. Once the contribution from these microbubbles is removed from the acoustic backscatter, the acoustic contrast of the nanobubbles was shown to be near noise levels. This result indicates that nanobubbles have limited utility as UCAs for CEUS.

## Introduction

Contrast-enhanced ultrasound (CEUS) is a safe, economical, portable and widely used medical imaging modality used in many clinical specialties, including cardiology,^1^ radiology^2^ and critical care^3^ medicine. Currently, lipid-coated, gas-filled microbubbles (MBs) (1-10 µm diameter) are clinically approved ultrasound contrast agents (UCAs) that provide a strong acoustic backscatter when imaged with clinical scanners operating at 1-10 MHz.^4^ When driven by an ultrasound pulse, the high compressibility of the gas within the MB causes the MB to oscillate volumetrically, which can be described by the modified Rayleigh-Plesset equation.^5,6^ MB acoustic scattering is much stronger than liquid or solid particles of similar size,^7^ thus providing strong contrast of the blood pool. MBs can therefore be used to measure blood flow^8^ and to image microvascular architecture.^9,10^ Researchers have also focused on bioconjugation methods, attaching ligands that can bind to endothelial receptors that serve as biomarkers for disease, such as tumor angiogenesis or inflammation.^11–13^

In clinical practice, MBs allow for high contrast images of tissues, such as kidneys, which are highly vascularized. However, in tissues with less vascularization, such as some solid tumors, MBs are less effective due to their confinement to the intravascular compartment.^14^ It has been shown that the amount of extravasation from leaky tumor vasculature via the enhanced permeability and retention (EPR) effect increases as particle size decreases.^15,16^ Therefore, decreasing the diameter of MBs into the nanoscale to have nanobubbles (NBs) (<1 µm diameter) should allow the NBs to extravasate into surrounding tissue, potentially enhancing tumor diagnosis and characterization via CEUS.^17^ However, recent studies have cast doubt on the utility of the EPR effect.^16^

Although NBs may extravasate to allow for extravascular imaging, experimental and modeling studies have shown that echogenicity decreases substantially with bubble size. For example, the linear model for ultrasound-driven bubble oscillations at 7 MHz predicts that the scattering cross section drops from 3.3 µm^2^ for a 1-µm radius bubble to 3.3×10^−6^ µm^2^ for a 100-nm radius bubble.^6,7^ Gorce et al. demonstrated experimentally and theoretically that a low threshold in diameter was found for SonoVue microbubbles at approximately 2 µm, under which size bubbles do not contribute appreciably to the echogenicity at medical ultrasound frequencies.^18^ Despite this prior work showing poor NB echogenicity, recent *in vivo* tumor CEUS imaging studies using NBs have concluded that NBs are echogenic.^19–23^

We hypothesize that this discrepancy can be explained by MB contamination in the NB suspensions. Prior CEUS studies have demonstrated that total bubble volume fraction is a more accurate prediction of *in vitro* and *in vivo* echogenicity than number concentration.^24,18^ Bubble volume scales with *R*^3^ (where *R* is bubble radius), so individual MBs contribute substantially more than NBs to the total bubble volume fraction. For example, a single 1-µm radius MB contributes the same gas volume as one-thousand 100-nm radius NBs. For a polydisperse mixture in which the vast majority of the number of particles are sub-micrometer in diameter, even a small number of MBs could account for a significant proportion of the total gas volume fraction. Therefore, in echogenicity studies of NBs, the echogenicity observed could be caused by the presence of small amounts of MBs.

The presence of a small amount of MBs may be missed by some of the sizing techniques that are used to characterize NBs. MBs are often sized by direct microscopy, electrozone sensing (EZS) or single particle optical sizing (SPOS).^25,26^ While these techniques ae suitable for MBs, they cannot detect NBs below the lower size threshold. Therefore, NBs are often characterized by light scattering (LS) or resonant mass measurement (RMM).^27–30^ However, these latter nanoparticle sizing techniques may miss the presence of MBs in the NB suspensions. Many of the NB studies have not reported results from MB sizing techniques, making it difficult to know whether there was MB contamination in those samples.

To test our hypothesis, we set out to determine the effect of MB contamination on NB+MB echogenicity, and to subtract this effect to determine more accurately the echogenicity of the NBs alone. We used multiple sizing techniques to characterize the size distribution the NB and MB formulations. To assess echogenicity, we used fundamental mode CEUS imaging at 7 MHz and 330 kPa peak negative pressure, which is consistent with prior studies that employed frequencies between 7 and 12 MHz and peak negative pressures between 200 and 500 kPa.^29,28,20,27,30^ Modeling simulations with the modified Rayleigh-Plesset equation were used to interpret our experimental results.

## Materials and Methods

### Reagents

1,2-dibehenoyl-sn-glycero-3-phosphocholine (DBPC) was purchased from Avanti Polar Lipids (Alabaster, AL). 1,2-dipalmitoyl-sn-glycero-3-phosphatidic acid (DPPA), 1,2-dipalmitoyl-sn-glycero-3-phosphoethanolamine (DPPE), and 1,2-distearoyl-sn-glycero-3-phosphoethanolamine-N-[amino(polyethylene glycol)-2000] (DSPE-PEG2000) were purchased from NOF America Corporation (White Plains, NY). Propylene glycol (PG) was purchased from Sigma-Aldrich (St. Louis, MO). Phosphate-buffered saline solution (PBS) was purchased from Fisher Scientific (Pittsburg, PA). Octafluoropropane (C_3_F_8_) was purchased from FluroMed (Round Rock, TX). The purity of all the reagents was ≥99%.

### Nanobubble Synthesis

The NB formulation was prepared in aqueous propylene glycol similar to the method described by de Leon et al.^28^ Briefly, stock solutions of 25 mL (10 mg/mL total lipid) were prepared comprising 0.61% (w/v) DBPC, 0.1% DPPA, 0.2% DPPE and 0.1% DSPE-PEG2000, dissolved in 20% (v/v) PG in PBS at 80 °C. The solution was then sonicated with a 20-kHz probe (Model 250A, Branson Ultrasonics; Danbury, CT) at low power (3/10; 3 W) for 10 min. 1 mL of this stock solution was then transferred to a 3-mL vial, the air inside the vial was replaced with C_3_F_8_ gas, and the vial was then mechanically agitated with a VialMix shaker (Lantheus, Billerica, MA) for 45 s. Up to 16 vials of NBs were prepared at a time, making approximately 20 mL of NB suspension stored in a 30-mL monoject syringe (Cardinal Health, Dublin, OH, USA).

### Size Selection of NBs

The size selection of the NB population were performed by differential centrifugation.^31,32^ The initial suspension was first centrifuged for 5 min at 50 RCF. The first 500 µL of the infranatant was collected into a 1-mL Eppendorf tube, and the rest of the infranatant was transferred to a 30-mL syringe, using a 16-G (1.194 mm inner diameter) needle. Then, 0.7 µm and 2.5 µm diameter NBs and MBs, respectively, were isolated. The samples were identified as crude (a), infranatant 5 min 50 RCF (b), first 500 µL infranatant 5 min 50 RCF (c), 0.7 size-selected NBs (d) and 2.5 size-selected MBs (e).

### Bubble Sizing

The number-weighted size distributions of the different samples were measured by light scattering (LS) using the Zetasizer Nano-S90 (Malvern, Worcestershire, UK), by single particle optical sizing (SPOS) using the AccuSizer 780A (Particle Sizing Systems, Santa Barbara, CA), or by electrozone sensing (EZS) using the Multisizer 3 (Coulter Counter, Beckman Coulter, Brea, CA). Samples of 0.5–150 µL were used for the AccuSizer and Multisizer, with the volume depending on sample concentration (largest volume for the least concentrated). For the Zetasizer, measurements were performed with 1:100 dilution of sample in filtered PBS loaded into a quartz cuvette. The mean particle diameters and concentrations were obtained from the number-weighted size distributions. The percentage of NBs (<1 µm) and MBs (1–2, 2–3, and >3 µm) using the EZS size distributions were also calculated from the number-weighted size distributions. Three independent preparations were evaluated for each formulation.

### Echogenicity

The CEUS echogenicity experiments were conducted using degassed 1.6% (w/v) agar phantoms in an agar mold with 16 G (1.65 mm outer diameter) sized channels, aligned roughly 2.5 cm from the top of the mold. The phantom was imaged using one-cycle, beam-focused (focus depth = 25 mm), fundamental mode pulse-echo imaging at 7 MHz, 330 kPa peak negative pressure (PNP) and 6 s pulse-repetition frequency (PRF) using a ULA-OP open ultrasound system (X-Phase, Florence, Italy) and an LA332 probe (Esaote SpA, Florence, Italy). The setup was imaged so that a centered cross section of the channel of interest was acquired for each experiment.

The bubble formulations described above (a-d) were diluted to various concentrations (10^3^ to 10^8^ bubbles/mL) with filtered PBS. The diluted sample was then transferred to a 3-mL monoject syringe (syringe diameter of 8.94 mm) fitted with a 16-G needle, which was then loaded onto an NE-300 infusion syringe pump (New Era Pump Systems, Farmingdale, NY) set to pump at 50 mL/h. A 1-mL sample was then collected after pumping, and its concentration was measured by EZS to estimate the number of particles that flowed through the channel during an experiment. The number of particles imaged was calculated by multiplying the particle concentration by the ultrasound beam volume (8.6×10^−3^ mL), which was calculated from the channel diameter and beam elevational width.

After priming the channel, 30 frames were taken sequentially under constant flow. Each frame was beamformed from the raw radiofrequency (RF) data. The pixel intensity was determined from the Hilbert transform of the RF data, and the pixel intensities from the 30 frames were averaged together. The contrast-to-noise ratio of each sample was calculated:

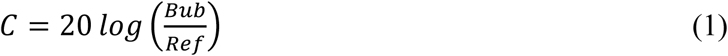

where *C* is the contrast-to-noise ratio in dB, *Bub* is the sum of all pixel intensities in the region of interest (ROI) of the channel containing the flowing MBs and NBs, and *Ref* is the sum of all pixel intensities in a region in the phantom of the same size as the *Bub* ROI at the same depth (a 5 mm × 5 mm^2^).

### Subtraction of the MB echoes

To remove the MB (>1 µm diameter) contribution, we used the non-log-compressed contrast enhancement (*CE*), defined by the following equation:

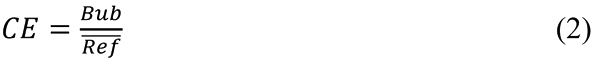

where 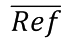 is average of all *Ref* values. We first calculated the *CE* from images obtained with the 2.5 µm size-isolated MB formulation and divided this value by the number of MBs greater than 2 µm. This calculation gave us the value of *CE*_*MB*>2*μm*_. Then, for each formulation, we determined the total *CE* for all bubbles less than 2 µm by the following equation to determine the percent contrast enhancement (%*CE*_*MBs*<2*μm*_):

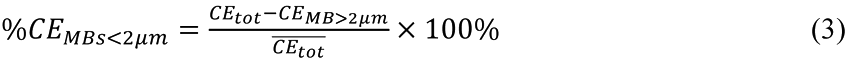

Finally, to determine the percent contrast enhancement due to NBs (%*CE*_*NBs*_), we plotted the %*CE*_*MBs*<2*μm*_ value as a function of the number of MBs with a diameter between 1 and 2 µm. We fit the data to a linear least-squares regression curve, and then extrapolated to the y-intercept (0%). This regression was reported with a 95% confidence interval.

### Simulating Ultrasonic Scattering from Bubbles

The predicted acoustic emission of NBs was determined by simulating the pressure scattered by a single bubble with a diameter ranging from 0.1 to 20 µm using the *BubbleSim* simulator in MATLAB by Lars Hoff.^6^ The *BubbleSim* model is limited in that it does not capture the full nonlinear rheology of the lipid coating, such as rupture and buckling captured by the Marmottant model^5^ for large-amplitude oscillations. However, the *BubbleSim* model does capture the nonlinear behavior of the gas core, which is expected to dominate the behavior of the bubble at this relatively high acoustic forcing at 7 MHz and 330 kPa PNP. The model ultrasound parameters were chosen to match the experimental pulse parameters used in the pulse-echo imaging study as closely as possible (frequency *f* = 7 MHz, peak negative pressure *PNP* = 330 kPa, number of cycles *nc* = 1, Hanning envelope). The bubble shell parameters were chosen to approximate the linear response of the lipid coating (shear modulus of the shell *G*_*s*_ = 50 MPa, shear viscosity of the shell *η*_*s*_ = 0.6 Pa-s, and shell thickness *δ* = 3 nm).^33^ The gas was treated isothermally, and the modeled medium was chosen as water. To determine the scattered pressure due to the bubble, the model was solved numerically using the stiff variable order ODE solver in *BubbleSim*. The theoretical acoustic emission was determined by calculating the maximum of the analytic signal of the simulated scattered pressure by the bubble. The total scattered pressure of each sample was calculated by assuming linear superposition and taking the weighted average based on the measured number of bubbles at each diameter and the model-predicted acoustic emission for that bubble size. The bubble concentrations used in this study were relatively low, so we neglected multi-bubble scattering. We did not account for the geometry of the bubbles in relation to the ultrasound transducer, nor did we account for attenuation in the intervening medium. The model predictions are therefore only estimates used to assess the relative importance of different bubble sizes.

### Statistical analysis

All data are expressed as the mean ± standard deviation. Differences between two experimental groups were assessed using a Students’ t-test, and one-way analysis of variance (ANOVA) was used for multiple comparisons. Data were evaluated using GraphPad software (San Diego, CA). A p-value ≤ 0.05 was used to indicate statistical significance.

## Results and Discussion

### Characterization of Bubble Size Distributions

The number-weighted size distributions of the different bubble samples were characterized by LS (Zetasizer), SPOS (AccuSizer) and EZS (Multisizer) (Fig. 1). The size distributions obtained for the crude bubble suspension showed a high composition of both NBs and MBs (Fig. 1a). Size-isolation by centrifugation progressively increased the fraction of NBs (<1 µm diameter, Fig. 1b-d) and decreased the fraction of MBs (>1 µm diameter, Fig. 1e).

**Figure 1.**
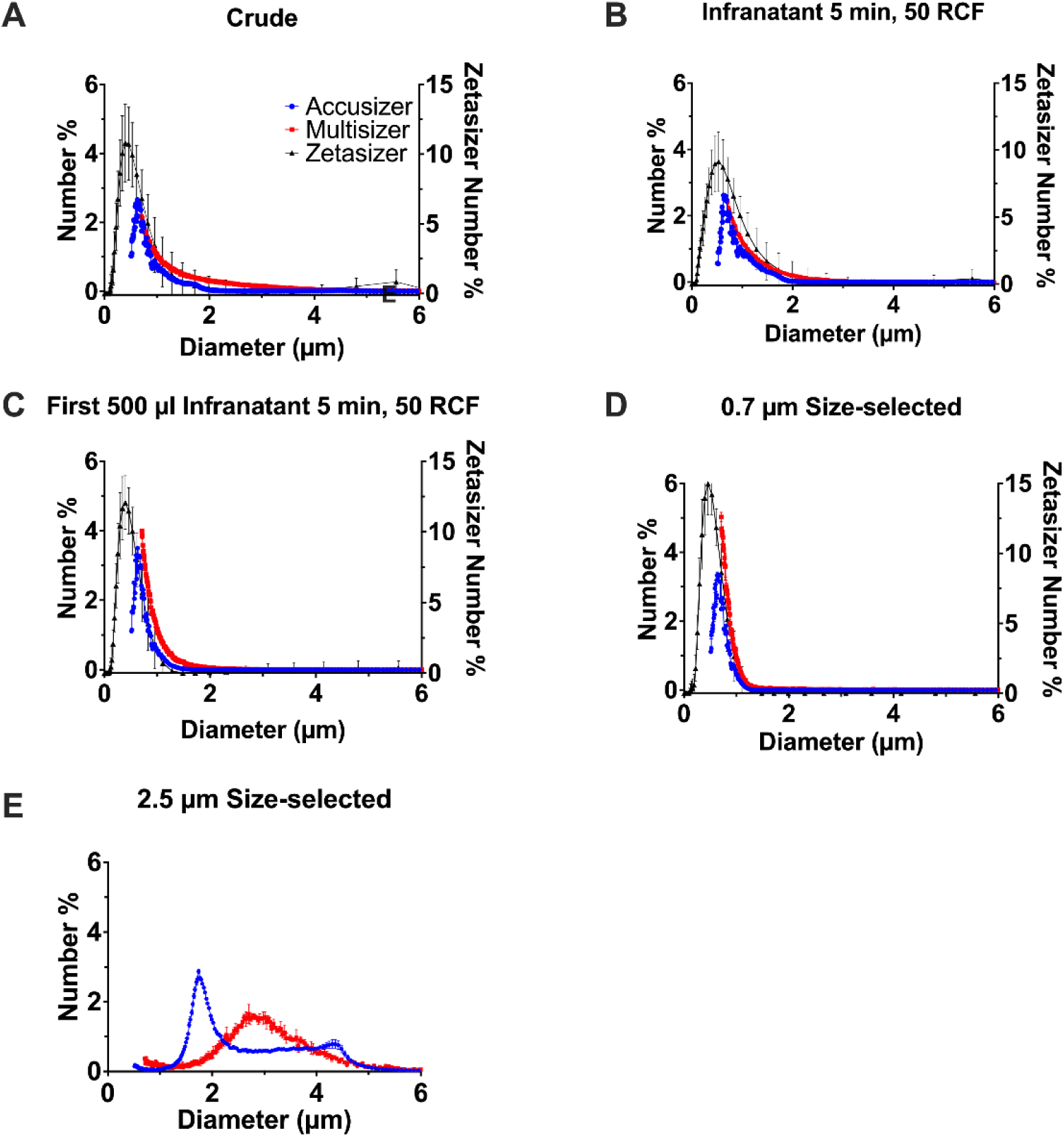
Number-weighted size distributions of NB and MB formulations measured using the LS (Zetasizer), SPOS (AccuSizer) and EZS (Multisizer). A) crude, prior to any size selection, B) infranatant after 5 min of centrifugation at 50 RCF, C) first 500 µL of the infranatant after 5 min of centrifugation at 50 RCF, D) after size selection to 0.7 µm, and E) after size selection to 2.5 µm MBs. Data represent the mean ± standard deviation for three independent preparations.

The number-weighted mean diameters obtained for each sample from LS (Zetasizer), SPOS (Accusizer) and EZS (Multisizer) are shown in Figure 2A. The number-weighted mean diameters obtained from LS, SPOS and EZS for the crude sample was 0.6±0.7, 0.9±0.6 and 1.3±0.9 µm, respectively (Fig. 2Aa). The crude sample could therefore be considered a NB formulation according to LS and SPOS. For the size-selected (0.7 µm size-selected) NBs, the number-weighted mean diameter was 0.5±0.2, 0.7±0.1 and 0.9±0.4, respectively (Fig. 2Ad). The 0.7 µm size-selected sample could therefore be considered a NB formulation according to all three sizing methods. For the 2.5 µm size-selected MBs, the number-weighted mean diameter obtained from SPOS and EZS was 2.4±1.0 and 2.7±1.0 µm (Fig. 2Ae), respectively (LS was not performed on this sample).

**Figure 2.**
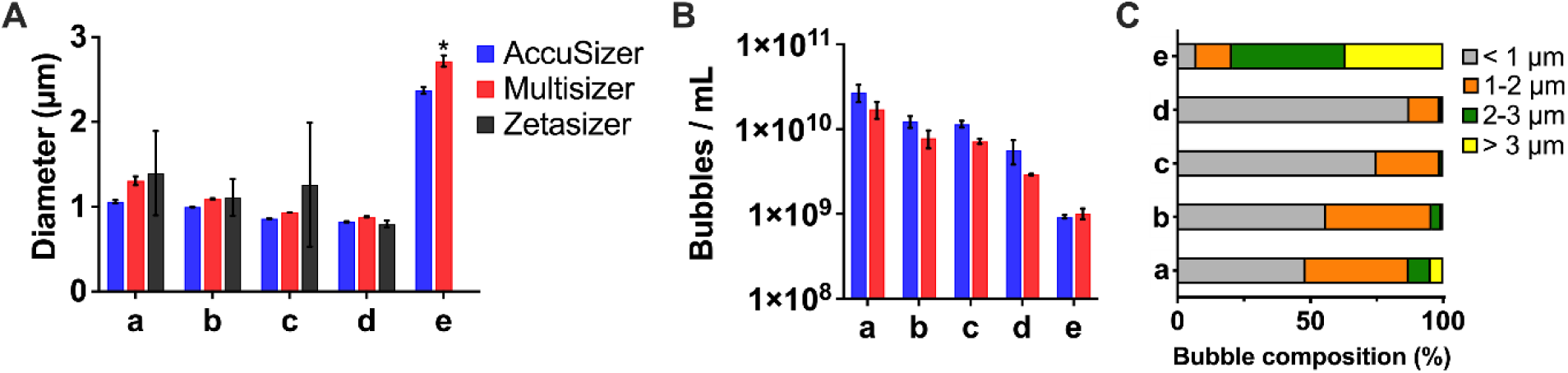
A) Number-weighted mean diameter of bubble formulations measured by SPOS (AccuSizer), EZS (Multisizer) and LS (Zetasizer). ^*^p < 0.05 *vs*. SPOS. B) Number-weighted mean bubble concentration obtained by SPOS and EZS. C) Proportion of bubbles <1 µm, 1–2 µm, 2–3 µm and > 3 µm detected by EZS. The bubble samples are labeled as crude (a), infranatant 5 min 50 RCF (b), first 500 µL infranatant 5 min 50 RCF (c), 0.7 µm size-selected NBs (d) and 2.5 µm size-selected MBs (e). Data represent the mean ± standard deviation for three independent preparations.

The concentration of each bubble sample was compared by using the SPOS and EZS methods (Fig. 2B). LS does not measure particle concentration. All the bubble preparation yielded reproducible bubble concentrations without statistical differences, ranging from 1×10^9^ to 5×10^10^ bubbles/mL. The trend is as expected, as sequential centrifugation steps lead to concomitant loss of MBs from the NB formulaitons.

The proportion of bubble diameters <1, 1**–**2, 2**–**3 and >3 µm present in the suspensions based on EZS measurements are also shown (Fig. 2C). The crude sample comprised 48% NBs (<1 µm), the infranatant 5 min, 50 RCF sample comprised 56% NBs, 40% 1**–**2 µm MBs and 4% >2 µm MBs. The first 500 µL infranatant 5 min, 50 RCF sample comprised 75% NBs, 24% 1**–**2 µm MBs and 1% >2 µm MBs. The 0.7 µm size-selected sample comprised 87% NBs and 13% >1µm MBs. Finally, the 2.5 µm size-isolated MB formulation comprised only 7% NBs and greater than 80% >2 µm MBs. The high fraction of MBs in the latter formulation allowed us to determine the echogenicity of >2 µm MBs and to remove this contribution from the total percent contrast enhancement of the NB formulations.

These results were used in the subsequent analysis of MB and NB echogenicity. Taken together, these data clearly show that NB formulations, at least those classified by a number-weighted mean diameter <1 µm, may contain a considerable number of MBs. This result suggests that multiple sizing techniques, including those that primarily size MBs, such as SPOS and EZS, should be used to characterize any NB formulation.

### Echogenicity of Bubble Formulations

The CEUS contrast for each bubble sample was calculated by analysis of ultrasound images of the agar phantom. Figure 3A shows an example of an ultrasound image, with the bright echoic region corresponding to a cross section of the channel through which the bubble sample was pumped. A region of interest was drawn around the channel (*Bub*) and in an adjacent part of the agar phantom (*Ref*), as shown.

**Figure 3.**
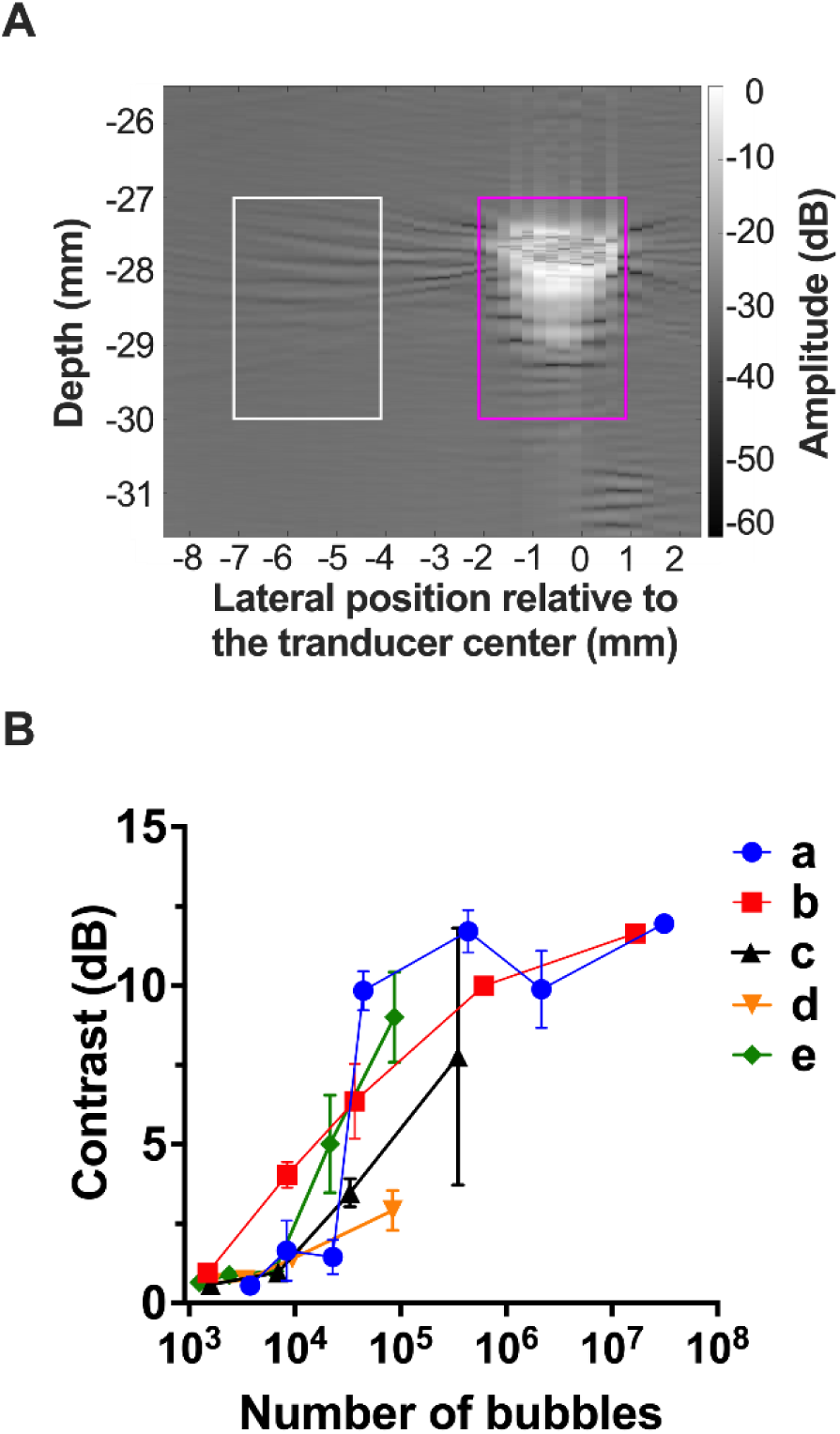
Contrast-to-noise ratios for the bubble formulations. A) Representative ultrasound contrast image, where the *Bub* region of interest is boxed in magenta and the *Ref* region is boxed in white. B) Contrast-to-noise ratio *vs*. particle number in the ultrasound beam volume. The bubble formulations were identified as crude (a), infranatant 5 min 50 RCF (b), first 500 µL infranatant 5 min 50 RCF (c), 0.7 µm size-selected NBs (d) and 2.5 µm size-selected MBs (e). ^*^p <0.05 *vs*. formulation (c). Data represent the mean ± standard deviation for three independent preparations.

The contrast-to-noise ratio calculated by equation (1) versus the number of bubbles excited within the beam volume is plotted in Figure 3B. In general, the size-selected formulations with the lowest proportions of MBs (Fig. 3Bc,d) produced the lower contrast than those comprising a greater fraction of MBs (Fig. 3Ba,b,e). For example, at 10^5^ particles, the formulation with the greatest NB fraction had a contrast-to-noise ratio of only 3 dB (Fig. 3d) compared to 10 dB for the crude formulation (Fig. 3Ba).

### Echogenicity of NBs

To compare echogenicity of bubbles less than 2 µm diameter in the bubble formulations, we used equation (3) to calculate the percent contrast enhancement (%*CE*_*MBs*<2*μm*_). These values are plotted in Figure 4A. It is interesting to note that only formulation (b) had a %*CE*_*MBs*<2*μm*_ that was statistically different than the crude formulation (which had a %*CE*_*MBs*<2*μm*_ ∼ 0%). This formulation had the greatest relative number of 1–2 µm MBs (Fig. 2c). We speculate that the echogenicity was relatively higher in this formulation owing to bubble resonance. The resonance frequency of ∼1.5 µm diameter MBs is ∼7 MHz.^34^ However, the totality of our data clearly show that echogenicity increases with increasing MB size.

**Figure 4.**
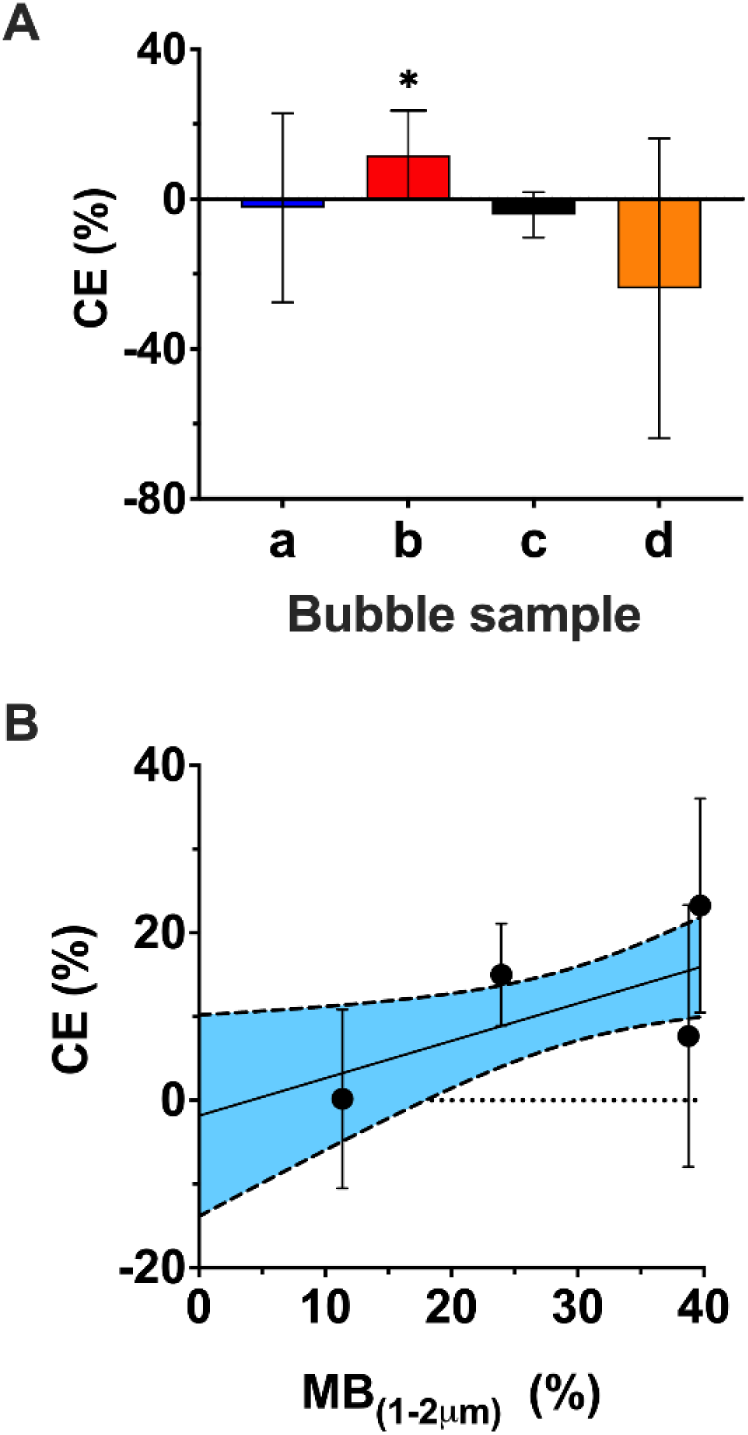
A) Percent contrast enhancement (%*CE*_*MBs*<2*μm*_) produced by bubbles <2 µm diameter for each formulation (a-e). The bubble formulations were identified as crude (a), infranatant 5 min 50 RCF (b), first 500 µL infranatant 5 min 50 RCF (c), 0.7 µm size-selected NBs (d) and 2.5 µm size-selected MBs (e). ^*^p <0.05 *vs*. formulation (a). B) Contribution to contrast enhancement from MBs between 1–2 µm diameter. A linear regression curve was fit to the data with a shaded 95% confidence interval extrapolated to a sample composed of 0% MBs. The y-intercept of the regression curve thus represents the contribution of NBs to the echogenicity. Data represent the mean ± standard deviation for three independent MB preparations.

Finally, the percent contrast enhancement (%*CE*_*NBs*_) associated to NBs (<1 µm diameter) was determined by plotting %*CE*_*MBs*<2*μm*_ versus number of MBs with a diameter between 1 and 2 µm (Fig. 4B). A linear least-squares regression showed a general increase in %*CE*_*MBs*<2*μm*_ with 1–2 µm MB number. The y-intercept of the regression curve is therefore an estimate of the percent contribution of only NBs (%*CE*_*NBs*_). From the 95% confidence interval, the contribution of NBs to contrast enhancement is only −1.84% *±* 12.00%. This result supports our hypothesis that MB contamination in NB formulations is responsible for the surprising echogenicity, at least under these ultrasound imaging conditions.

### Theoretical Scattered Pressure of NBs

The maximum scattered pressure from a bubble excited by an ultrasound pulse of the same parameters used experimentally was simulated by using the *BubbleSim* software.^6^ The scattered pressure was simulated for bubble diameters ranging from 0.1 to 20 µm (Fig. 5A). Notably, the simulated scattered pressure increases from ∼10^−5^ Pa for a 100-nm zdiameter bubble to ∼10^3^ Pa for a 1-µm diameter bubble. This million-fold increase in echogenicity is consistent with the scattering cross section estimation from the linear model for bubble oscillations (e.g., from 3.3×10^−6^ µm^2^ for a 100-nm radius bubble to 3.3 µm^2^ for a 1-µm radius bubble at 7 MHz).^6,7^ According to the *BubbleSim* model, a single 0.7-µm diameter NB scatters only 2.6% of that for a 2.5-µm diameter MB. Figure 5B shows the predicted total scattered pressure for the ensemble of bubbles *vs*. bubble number for each of the formulations. The predictions are consistent with the general trend for the observed contrast enhancement obtained experimentally from each bubble formulation, as plotted in Fig. 3B. Thus, the modeling results support our hypothesis that a few contaminating MBs may contribute disproportionally to the surprising echogenicity of NB formulations. The qualitative differences between the curves in Fig. 3B and Fig. 5B is likely due to the model assumptions, such as neglecting the relative bubble/transducer geometry and attenuation (discussed further below).

**Figure 5.**
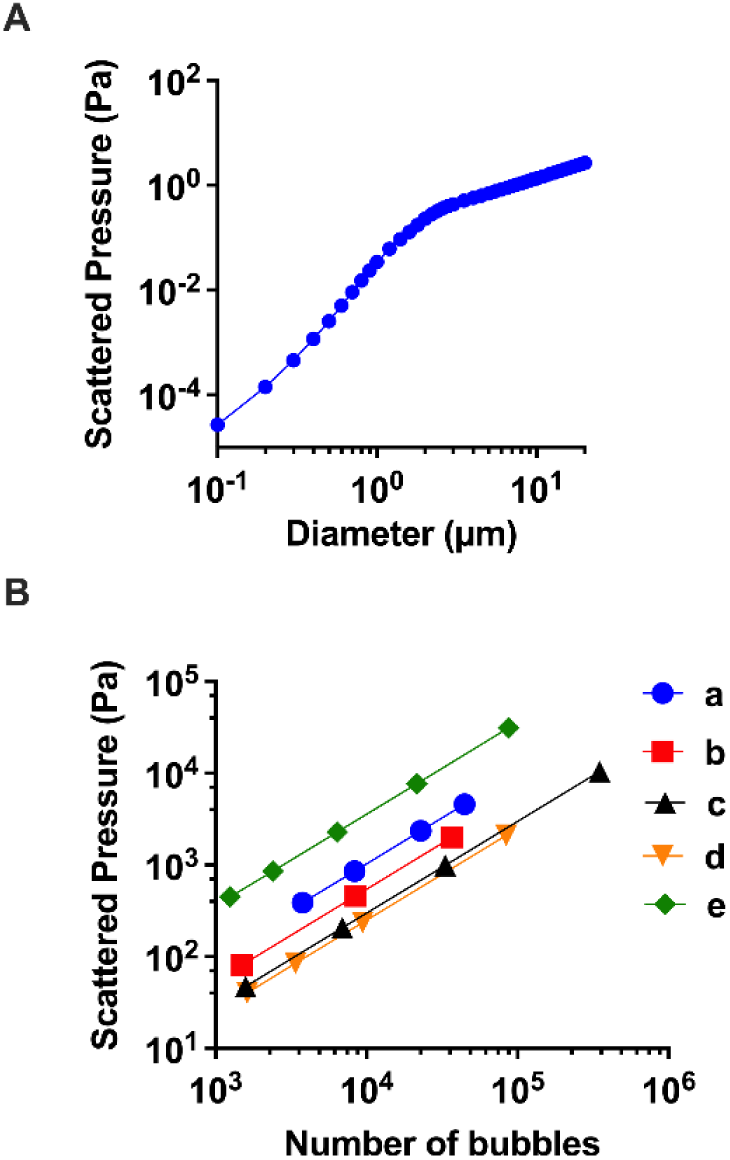
A) Simulated scattered pressure from a single bubble by solving a modified form of the Rayleigh-Plesset equation using *BubbleSim*. The bubble was irradiated at 7 MHz, 330 kPa peak negative pressure, single cycle and Hanning envelope. Simulation numerically solved B) Simulated amount of pressure that each NB formulation sample (a-e) would scatter versus the number of irradiated bubbles in the sample. The bubble samples were identified as crude (a), infranatant 5 min 50 RCF (b), first 500 µL infranatant 5 min 50 RCF (c), 0.7 µm size-selected NBs (d) and 2.5 µm size-selected MBs (e).

## Limitations of this Study

The main limitations of this study are as follows: 1) We only examined fundamental-mode imaging at one frequency (7 MHz) and one acoustic pressure (330 kPa PNP). Therefore, we did not fully explore the full acoustic spectrum of the bubble response, such as sub-harmonics and ultra-harmonics, nor did we examine nonlinear effects that can be exploited in amplitude-modulation, pulse-inversion and other nonlinear CEUS imaging modes. 2) We only examined one lipid/gas formulation. It is conceivable that other shells or gases could impact the bubble echogenicity under these ultrasound conditions. 3) The *BubbleSim* model was a rough estimate of the theoretical bubble response. A more sophisticated model, such as the Chatterjee-Sarkar^**35**^ model or the Marmottant^**5**^ model for the lipid shell viscoelasticity could be used. Additionally, the modeling could be improved by including effects of microbubble instability, the relative bubble/transducer geometry, frequency-dependent acoustic attenuation, and other experimental factors.

## Conclusion

In this study, we examined the echogenicity of various formulations of DBPC:DSPE-PEG2000 shelled C_3_F_8_ gas filled microbubbles with different size distributions. The formulations were characterized by three different sizing methods, and MBs (>1 µm diameter) were found in all formulations. This result suggests that any NB (<1 µm diameter) formulation should be evaluated for size distribution with multiple methods, especially those specific to MBs, such as single particle optical sensing or electrozone sensing. Additionally, our pulse-echo imaging analysis of the different formulations indicated that echogenicity is due to the MBs, and that NBs contribute essentially nothing to the total acoustic backscatter. This observation was supported by modeling. Taken together, this study supports our hypothesis that contaminating MBs are responsible for the echogenicity of NB formulations. Limitations to this study include the use of only one pulse-echo imaging mode, the use of only one bubble formulation chemistry, and simplifying assumptions in the modeling.

## Acknowledgements

This research was funded by NIH grant R01CA195051 to MB.

## Graphical TOC Entry

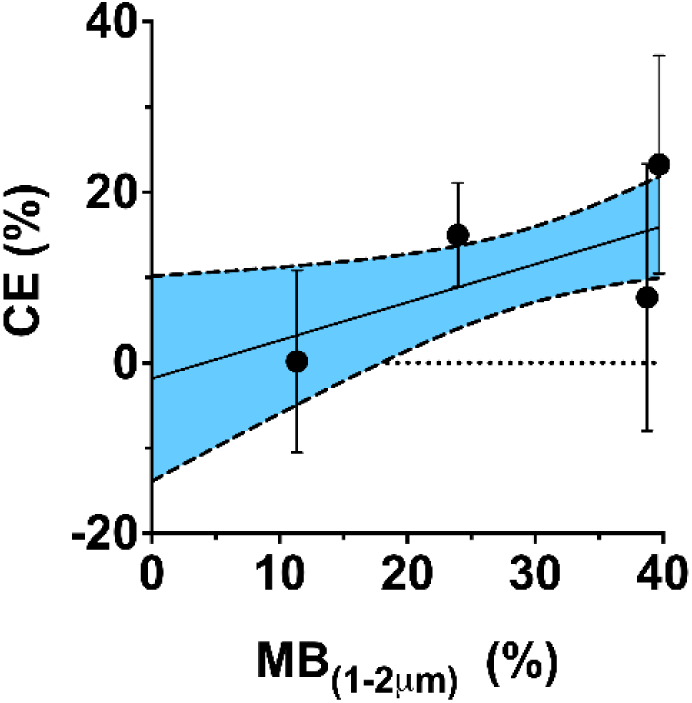

## References

(1) Lindner, J. R. Contrast Echocardiography: Current Status and Future Directions. Heart 2021, 107 (1), 18–24. https://doi.org/10.1136/heartjnl-2020-316662.

(2) Wilson, S. R.; Burns, P. N.; Kono, Y. Contrast-Enhanced Ultrasound of Focal Liver Masses: A Success Story. Ultrasound in Medicine & Biology 2020, 46 (5), 1059–1070. https://doi.org/10.1016/j.ultrasmedbio.2019.12.021.

(3) Kummer, T.; Oh, L.; Phelan, M. B.; Huang, R. D.; Nomura, J. T.; Adhikari, S. Emergency and Critical Care Applications for Contrast-Enhanced Ultrasound. The American Journal of Emergency Medicine 2018, 36 (7), 1287–1294. https://doi.org/10.1016/j.ajem.2018.04.044.

(4) Stride, E.; Segers, T.; Lajoinie, G.; Cherkaoui, S.; Bettinger, T.; Versluis, M.; Borden, M. Microbubble Agents: New Directions. Ultrasound in Medicine & Biology 2020, 46 (6), 1326–1343. https://doi.org/10.1016/j.ultrasmedbio.2020.01.027.

(5) Marmottant, P.; van der Meer, S.; Emmer, M.; Versluis, M.; de Jong, N.; Hilgenfeldt, S.; Lohse, D. A Model for Large Amplitude Oscillations of Coated Bubbles Accounting for Buckling and Rupture. The Journal of the Acoustical Society of America 2005, 118 (6), 3499–3505. https://doi.org/10.1121/1.2109427.

(6) Hoff, L. Acoustic Characterization of Contrast Agents for Medical Ultrasound Imaging; Springer Science & Business Media, 2013.

(7) Borden, M. A.; Song, K.-H. Reverse Engineering the Ultrasound Contrast Agent. Adv Colloid Interface Sci 2018, 262, 39–49. https://doi.org/10.1016/j.cis.2018.10.004.

(8) Reisner, S. A.; Ong, L. S.; Lichtenberg, G. S.; Amico, A. F.; Shapiro, J. R.; Allen, M. N.; Meltzer, R. S. Myocardial Perfusion Imaging by Contrast Echocardiography with Use of Intracoronary Sonicated Albumin in Humans. J Am Coll Cardiol 1989, 14 (3), 660–665. https://doi.org/10.1016/0735-1097(89)90107-1.

(9) Gessner, R. C.; Aylward, S. R.; Dayton, P. A. Mapping Microvasculature with Acoustic Angiography Yields Quantifiable Differences between Healthy and Tumor-Bearing Tissue Volumes in a Rodent Model. Radiology 2012, 264 (3), 733–740. https://doi.org/10.1148/radiol.12112000.

(10) Errico, C.; Pierre, J.; Pezet, S.; Desailly, Y.; Lenkei, Z.; Couture, O.; Tanter, M. Ultrafast Ultrasound Localization Microscopy for Deep Super-Resolution Vascular Imaging. Nature 2015, 527 (7579), 499–502. https://doi.org/10.1038/nature16066.

(11) Klibanov, A. L. Ligand-Carrying Gas-Filled Microbubbles: Ultrasound Contrast Agents for Targeted Molecular Imaging. Bioconjugate Chem. 2005, 16 (1), 9–17. https://doi.org/10.1021/bc049898y.

(12) Pillai, R.; Marinelli, E. R.; Fan, H.; Nanjappan, P.; Song, B.; von Wronski, M. A.; Cherkaoui, S.; Tardy, I.; Pochon, S.; Schneider, M.; Nunn, A. D.; Swenson, R. E. A Phospholipid−PEG2000 Conjugate of a Vascular Endothelial Growth Factor Receptor 2 (VEGFR2)-Targeting Heterodimer Peptide for Contrast-Enhanced Ultrasound Imaging of Angiogenesis. Bioconjugate Chem. 2010, 21 (3), 556–562. https://doi.org/10.1021/bc9005688.

(13) Slagle, C. J.; Thamm, D. H.; Randall, E. K.; Borden, M. A. Click Conjugation of Cloaked Peptide Ligands to Microbubbles. Bioconjugate Chem. 2018, 29 (5), 1534–1543. https://doi.org/10.1021/acs.bioconjchem.8b00084.

(14) Moestue, S. A.; Gribbestad, I. S.; Hansen, R. Intravascular Targets for Molecular Contrast-Enhanced Ultrasound Imaging. IJMS 2012, 13 (6), 6679–6697. https://doi.org/10.3390/ijms13066679.

(15) Ishida, O.; Maruyama, K.; Sasaki, K.; Iwatsuru, M. Size-Dependent Extravasation and Interstitial Localization of Polyethyleneglycol Liposomes in Solid Tumor-Bearing Mice. International Journal of Pharmaceutics 1999, 190 (1), 49–56. https://doi.org/10.1016/S0378-5173(99)00256-2.

(16) Izci, M.; Maksoudian, C.; Manshian, B. B.; Soenen, S. J. The Use of Alternative Strategies for Enhanced Nanoparticle Delivery to Solid Tumors. Chem. Rev. 2021, 121 (3), 1746–1803. https://doi.org/10.1021/acs.chemrev.0c00779.

(17) Duan, L.; Yang, L.; Jin, J.; Yang, F.; Liu, D.; Hu, K.; Wang, Q.; Yue, Y.; Gu, N. Micro/Nano-Bubble-Assisted Ultrasound to Enhance the EPR Effect and Potential Theranostic Applications. Theranostics 2020, 10 (2), 462–483. https://doi.org/10.7150/thno.37593.

(18) Gorce, J.-M.; Arditi, M.; Schneider, M. Influence of Bubble Size Distribution on the Echogenicity of Ultrasound Contrast Agents: A Study of SonoVue??? Investigative Radiology 2000, 35 (11), 661–671. https://doi.org/10.1097/00004424-200011000-00003.

(19) Xing, Z.; Wang, J.; Ke, H.; Zhao, B.; Yue, X.; Dai, Z.; Liu, J. The Fabrication of Novel Nanobubble Ultrasound Contrast Agent for Potential Tumor Imaging. Nanotechnology 2010, 21 (14), 145607. https://doi.org/10.1088/0957-4484/21/14/145607.

(20) Pellow, C.; Abenojar, E. C.; Exner, A. A.; Zheng, G.; Goertz, D. E. Concurrent Visual and Acoustic Tracking of Passive and Active Delivery of Nanobubbles to Tumors. Theranostics 2020, 10 (25), 11690–11706. https://doi.org/10.7150/thno.51316.

(21) Yu, Z.; Wang, Y.; Xu, D.; Zhu, L.; Hu, M.; Liu, Q.; Lan, W.; Jiang, J.; Wang, L. G250 Antigen-Targeting Drug-Loaded Nanobubbles Combined with Ultrasound Targeted Nanobubble Destruction: A Potential Novel Treatment for Renal Cell Carcinoma. IJN 2020, Volume 15, 81–95. https://doi.org/10.2147/IJN.S230879.

(22) Zhou, T.; Cai, W.; Yang, H.; Zhang, H.; Hao, M.; Yuan, L.; Liu, J.; Zhang, L.; Yang, Y.; Liu, X.; Deng, J.; Zhao, P.; Yang, G.; Duan, Y. Annexin V Conjugated Nanobubbles: A Novel Ultrasound Contrast Agent for in Vivo Assessment of the Apoptotic Response in Cancer Therapy. Journal of Controlled Release 2018, 276, 113–124. https://doi.org/10.1016/j.jconrel.2018.03.008.

(23) Cai, W. B.; Yang, H. L.; Zhang, J.; Yin, J. K.; Yang, Y. L.; Yuan, L. J.; Zhang, L.; Duan, Y. Y. The Optimized Fabrication of Nanobubbles as Ultrasound Contrast Agents for Tumor Imaging. Sci Rep 2015, 5 (1), 13725. https://doi.org/10.1038/srep13725.

(24) Sontum, P. C.; Ostensen, J.; Dyrstad, K.; Hoff, L. Acoustic Properties of NC100100 and Their Relation with the Microbubble Size Distribution. Invest Radiol 1999, 34 (4), 268–275. https://doi.org/10.1097/00004424-199904000-00003.

(25) Sennoga, C. A.; Yeh, J. S. M.; Alter, J.; Stride, E.; Nihoyannopoulos, P.; Seddon, J. M.; Haskard, D. O.; Hajnal, J. V.; Tang, M.-X.; Eckersley, R. J. Evaluation of Methods for Sizing and Counting of Ultrasound Contrast Agents. Ultrasound in Medicine & Biology 2012, 38 (5), 834–845. https://doi.org/10.1016/j.ultrasmedbio.2012.01.012.

(26) Satinover, S. J.; Dove, J. D.; Borden, M. A. Single-Particle Optical Sizing of Microbubbles. Ultrasound in Medicine & Biology 2014, 40 (1), 138–147. https://doi.org/10.1016/j.ultrasmedbio.2013.08.018.

(27) Abenojar, E. C.; Bederman, I.; de Leon, A. C.; Zhu, J.; Hadley, J.; Kolios, M. C.; Exner, A. Theoretical and Experimental Gas Volume Quantification of Micro- and Nanobubble Ultrasound Contrast Agents. Pharmaceutics 2020, 12 (3), 208. https://doi.org/10.3390/pharmaceutics12030208.

(28) de Leon, A.; Perera, R.; Hernandez, C.; Cooley, M.; Jung, O.; Jeganathan, S.; Abenojar, E.; Fishbein, G.; Sojahrood, A. J.; Emerson, C. C.; Stewart, P. L.; Kolios, M. C.; Exner, A. A. Contrast Enhanced Ultrasound Imaging by Nature-Inspired Ultrastable Echogenic Nanobubbles. Nanoscale 2019, 11 (33), 15647–15658. https://doi.org/10.1039/C9NR04828F.

(29) Abenojar, E. C.; Nittayacharn, P.; de Leon, A. C.; Perera, R.; Wang, Y.; Bederman, I.; Exner, A. A. Effect of Bubble Concentration on the in Vitro and in Vivo Performance of Highly Stable Lipid Shell-Stabilized Micro- and Nanoscale Ultrasound Contrast Agents. Langmuir 2019, 35 (31), 10192–10202. https://doi.org/10.1021/acs.langmuir.9b00462.

(30) Jafari Sojahrood, A.; de Leon, A. C.; Lee, R.; Cooley, M.; Abenojar, E. C.; Kolios, M. C.; Exner, A. A. Toward Precisely Controllable Acoustic Response of Shell-Stabilized Nanobubbles: High Yield and Narrow Dispersity. ACS Nano 2021, 15 (3), 4901–4915. https://doi.org/10.1021/acsnano.0c09701.

(31) Sirsi, S.; Feshitan, J.; Kwan, J.; Homma, S.; Borden, M. Effect of Microbubble Size on Fundamental Mode High Frequency Ultrasound Imaging in Mice. Ultrasound in Medicine & Biology 2010, 36 (6), 935–948. https://doi.org/10.1016/j.ultrasmedbio.2010.03.015.

(32) Feshitan, J. A.; Chen, C. C.; Kwan, J. J.; Borden, M. A. Microbubble Size Isolation by Differential Centrifugation. Journal of Colloid and Interface Science 2009, 329 (2), 316– 324. https://doi.org/10.1016/j.jcis.2008.09.066.

(33) Tu, J.; Guan, J.; Qiu, Y.; Matula, T. J. Estimating the Shell Parameters of SonoVue ® Microbubbles Using Light Scattering. The Journal of the Acoustical Society of America 2009, 126 (6), 2954–2962. https://doi.org/10.1121/1.3242346.

(34) Doinikov, A. A.; Haac, J. F.; Dayton, P. A. Resonance Frequencies of Lipid-Shelled Microbubbles in the Regime of Nonlinear Oscillations. Ultrasonics 2009, 49 (2), 263–268. https://doi.org/10.1016/j.ultras.2008.09.006.

(35) Chatterjee, D.; Sarkar, K. A Newtonian Rheological Model for the Interface of Microbubble Contrast Agents. Ultrasound Med Biol 2003, 29 (12), 1749–1757. https://doi.org/10.1016/s0301-5629(03)01051-2.

